# Behavioral interactions between bacterivorous nematodes and predatory bacteria in a synthetic community

**DOI:** 10.1101/2021.04.20.440615

**Authors:** Nicola Mayrhofer, Gregory J. Velicer, Kaitlin A. Schaal, Marie Vasse

## Abstract

Theory and empirical studies in metazoans predict that apex predators should shape the behavior and ecology of mesopredators and prey at lower trophic levels. Despite the ecological importance of microbial communities, few studies of predatory microbes examine such behavioral responses and the multiplicity of trophic interactions. Here, we sought to assemble a three-level microbial food chain and to test for behavioral interactions between the predatory nematode *Caenorhabditis elegans* and the predatory social bacterium *Myxococcus xanthus* when cultured together with two basal prey bacteria that both predators can eat - *Escherichia coli* and *Flavobacterium johnsoniae*. We find that >90% of *C. elegans* worms failed to interact with *M. xanthus* even when it was the only potential prey species available, whereas most worms were attracted to pure patches of *E. coli* and *F. johnsoniae*. In addition, *M. xanthus* altered nematode predatory behavior on basal prey, repelling *C. elegans* from two-species patches that would be attractive without *M. xanthus*, an effect similar to that of *C. elegans* pathogens. The nematode also influenced the behavior of the bacterial predator: *M. xanthus* increased its predatory swarming rate in response to *C. elegans* in a manner dependent both on basal-prey identity and on worm density. Our results suggest that *M. xanthus* is an unattractive prey for some soil nematodes and is actively avoided when more lucrative prey are available. Most broadly, we find that nematode and bacterial predators mutually influence one another’s predatory behavior, with likely consequences for coevolution within complex microbial food webs.

## Introduction

Predation is an ancient biological interaction that influences ecosystem resource turnover^1,2^ as well as species abundance, diversity, and evolution^3–8^. Predators can be found at all biological scales and include organisms as different as white sharks and microbes. While less familiar, small predators such as protists, nematodes, and bacteria make fundamental contributions to global biogeochemical cycling^9,10^ and are proposed to be key players for both agriculture^11^ and human health^12,13^. In addition, predation in microbial communities may have been a driving force in some of the major transitions in evolution, including the origin of the eukaryotic cell^14–17^ and the advent of multicellularity^18,19^.

The predatory interactions that link members of a community can be represented as food webs, trophic networks that display the flow of energy among community members. Food webs have long been a central concept in ecology and are powerful tools for investigating community structure, the nature and strength of pairwise interactions, and the indirect effects of interactions on various aspects of community ecology. Food web research typically relies on direct observation of organism behavior, but such direct observations are difficult or impossible when studying microbes (especially given that many microbes cannot be cultured under laboratory conditions). To study species interactions within microbial food webs^20^, ecologists rely on computational studies, mathematical modelling, and experiments in simplified microbial systems. Examples include flux-balance analysis^21–23^, study of pairwise interactions^24^, and examination of growth and death curves in small communities^25^. These approaches usually adopt a bottom-up strategy, inferring features of the community based on the careful investigation of its components. However, they likely miss out on higher-order interactions and complex behavioral responses. Since predatory interactions may occur in complex networks, involve various partners, and fluctuate over time, these bottom-up approaches might be insufficient to understand the dynamics and broader ecological impacts of microbial predation^26,27 25,28^.

Despite the ecological importance of microbes^10^, microbial predators have only recently received substantial recognition as agents that influence biodiversity by controlling and shaping bacterial communities^25,29–34^. Microbial predators use a wide range of strategies to kill and consume their prey, from the periplasm-invasion strategy of *Bdellovibrio bacteriovorus*, which grows and divides within its prey, to total engulfment by protists and far-range killing by *Streptomyces* species^35^. Cells of *Myxococcus xanthus* forage in groups, repeatedly reversing direction while attacking and consuming prey^36^. According to current understanding, *M. xanthus* secretes extracellular hydrolytic enzymes that break down prey macromolecules and allow uptake of the released nutrients^37^. Studies of *M. xanthus* predation have examined its molecular mechanisms^36,38,39^, the effects of ecological conditions^40–43^, and co-evolution with a single prey species^44,45^. Little is understood about how *M. xanthus* may itself be exposed to predation pressure and how it interacts with its own predators. Natural bacteria communities are often grazed by bacteriophagous microfauna such as nematodes and protozoa, which can influence their structure and composition^46,47^. It is very likely that some such bacteriophagous organisms prey upon *M. xanthus* in natural environments, thus making *M. xanthus* a mesopredator. In fact, Dahl *et al*.^48^ have shown that the predatory nematode *Caenorhabditis elegans* will ingest *M. xanthus* in some contexts. But whether *C. elegans* achieves net growth from nutrients derived from wild-type *M. xanthus* remains uncertain.

It is not known to what extent the community-ecology effects of microbial predators mirror those of multicellular predators. In large organisms, intraguild predation (when apex predator and mesopredator also compete for the same basal prey organism) can have direct effects on mesopredator survival and distribution^49^. Ritchie and Johnson^50^ reviewed the effects of apex predators on mesopredators and their prey in 94 animal studies and found that, on average, increasing apex predator population size two-fold reduces mesopredator abundance by approximately four-fold. Such effects may be similarly important in communities of microbes. In addition to their direct demographic effects, apex predators can generate substantial behavioral modifications in mesopredators, altering their habitat use and changing their foraging activity, thereby indirectly affecting their survival and growth^51^. It is unclear whether these communities show hierarchical trophic interactions, and if so whether bacteriophagous organisms function as apex predators. The study of microbial community dynamics therefore requires a better understanding of the direct and indirect interactions between bacterial predators (potential mesopredators) and bacteriophagous nematodes and protozoa (potential apex predators).

Here, we examined behavioral interactions between two predator species in a simple community hypothesized to involve a food chain with three trophic levels: *C. elegans* as a potential apex predator, *M. xanthus* as a potential mesopredator, and two basal prey bacteria (*E. coli* and *F. johnsoniae*). We first tested for effects of the bacterial predator on nematode predatory behavior by asking whether i) *M. xanthus* attracts or repels *C. elegans* in the absence of other prey, ii) bacterial cell death or strain motility alters any effect of *M. xanthus* on *C. elegans*, iii) *M. xanthus* is more or less attractive to *C. elegans* as potential prey than the two basal prey species, and iv) the presence of *M. xanthus* in mixture with one basal prey species in a given prey patch alters its attractiveness to worms. We then asked whether nematodes reciprocally influence *M. xanthus* behavior - specifically swarming behavior within patches of basal prey - whether due to direct interactions between the predator species or indirectly due to nematode effects on basal prey populations.

## Materials and Methods

### Bacterial and nematode strains

As the hypothesized apex predator, we used *C. elegans* strain N2 (CGC). As hypothesized mesopredators, we used two strains of *Myxococcus xanthus*, GJV1 and GJV71. *M. xanthus* uses two distinct motility systems to drive swarming across solid surfaces, traditionally referred to as the ‘A motility system’ and the ‘S motility system’^52^. Strain GJV1, aka strain ‘S’ in Velicer *et al*.^53^, possesses both systems functionally intact. For purposes of this paper, we hereafter refer to GJV1 as strain S, for ‘swarming’. GJV71 is a nonmotile mutant of GJV1 with major deletions in two genes, one gene essential for A motility (*cglB*) and one gene essential for S motility (*pilA*). GJV71 was referred to as strain ‘A1 *cglB*’ in Velicer & Yu^54^. We hereafter refer to GJV71 as strain N, for ‘non-swarming’. We selected the *Escherichia coli* strain OP50^55^ (CGC, Caenorhabditis Genetic Center) and *Flavobacterium johnsoniae* (ATCC® 17061™) as basal prey bacteria because they represent, respectively, high and intermediate quality food sources for *M. xanthus*, promoting *M. xanthus* swarming and growth to different degrees^42,43^. *E. coli* strain OP50 is the standard prey for laboratory populations of *C. elegans*^56^. In preliminary experiments, *F. johnsoniae* sometimes displayed a phenotype with low gliding ability which inhibited *M. xanthus* predation. In all following experiments, the source of *F. johnsoniae* was a frozen stock originating from a single colony which we isolated from a normally spreading population.

### Standard culture conditions

Unless otherwise indicated, organisms were cultured on 6-cm diameter petri dishes each with 14 ml of 1.5% agar CFcc medium (‘clone fruiting’ medium^57^ supplemented with 1 mM CaCl_2_ and 0.005 mg/ml cholesterol).

### Culturing C. elegans

We froze *C. elegans* in a 10% DMSO solution and thawed it in minimal salts buffer (M9) with glutamine, according to Pires da Silve *et al*.^58^. We maintained the worms at room temperature on 1.5% agar nematode growth medium (NGM) dishes seeded with *E. coli* OP50, transferring weekly. We synchronized the life stages of all *C. elegans* populations prior to use in an experiment. Seven days before the start of the experiment, we transferred a small inoculum from a growing population to seeded 1.5% agar high growth medium (HGM) dishes^59^. After six days of incubation at room temperature, we washed the agar surface with M9 to collect the worms. We centrifuged them at 173 x *g* for 1 minute and removed all but 1 ml of supernatant. We then added 3 ml of bleaching solution (6 ml ddH_2_O, 6 ml NaOCl (5% Cl), 2 ml 1M NaOH) and waited up to 6 minutes, vortexing every 2 minutes. This step dissolves the bodies of the adult worms, releasing the eggs. We then washed the released eggs four times by centrifuging, removing all but 500 µl of supernatant, and adding ddH_2_O to 5 ml. After the final wash, we added 3.5 ml of M9 and transferred the egg suspension to a 6-cm petri dish to hatch at room temperature overnight. To prevent contamination, we added 40 µg/ml gentamicin. The next day, we collected the hatched L1 larvae by centrifuging and resuspended them in CFcc liquid. We determined the worm density in the suspension by plating three 1-µl drops on unseeded CFcc agar plates and counting the worms in each drop.

### Culturing M. xanthus

We inoculated *M. xanthus* from freezer stock onto 1.5% agar CTT (10 g/l Casitone, 10 mM Tris pH 8.0, 8 mM MgSO_4_, 1 mM KPO_4_^60^) dishes and incubated it at 32 °C and 90% relative humidity (rH) for 4-5 days. We then sampled the outer edge of the resulting colony and transferred the inoculum into CTT liquid, shaking at 32 °C and 300 rpm for 1 day until the cultures reached mid-exponential phase, then adjusted them to an absorbance (OD_600_) of 5 in CFcc liquid.

### Culturing prey bacteria

We streaked *F. johnsoniae* and *E. coli* from freezer stock onto 1.5% agar lysogeny broth (LB, Sigma) dishes and incubated them at 32 °C and 90% rH for 3 days. We transferred single colonies into LB liquid, shaking at 32 °C and 300 rpm for 1 day (or ∼10 hours in the case of *E. coli*), then adjusted the cultures to an OD_600_ of 5 in CFcc liquid.

### C. elegans binary choice assays

We inoculated two 15-µl bacteria spots 2 cm apart on CFcc agar and incubated at 25 °C and 50% rH overnight before adding *C. elegans*. We added 20 *C. elegans* L1 larvae suspended in CFcc liquid to the dishes and incubated them at 25 °C and 50% rH. We counted the number of worms in different regions of the petri dish under a dissecting microscope at several time points. For the assays reported in Fig. 1A,B, we prepared both living and dead cultures of *M. xanthus* strains S and N. To kill *M. xanthus*, we resuspended a growing culture to OD 5 in CFcc liquid and incubated at 50 °C for 3 hours. We left live cells shaking at 32 °C and 300 rpm during this time. Results from these assays are shown in Fig. 1A,B, Fig. 2, and Fig. 3.

**Figure 1.**
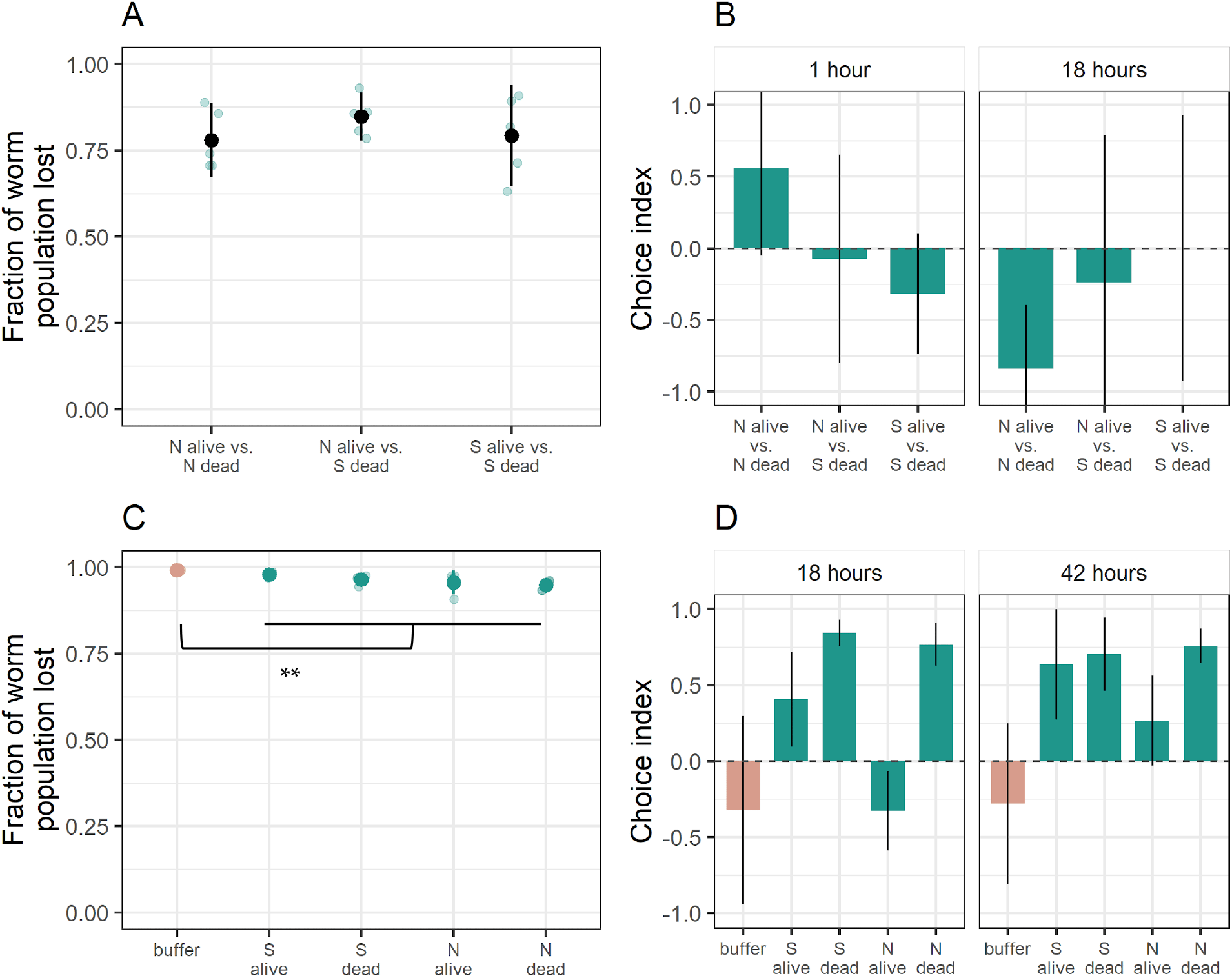
*C. elegans* prefers *M. xanthus* to buffered agar. We tested potential effects of living or dead *M. xanthus* on the position of worms presented with choices between either two circular patches of *M. xanthus* (A, B) or between a half-plate lawn of *M. xanthus* versus a half plate of bacteria-free agar (C, D). For both experiments, we report the mean fraction of *C. elegans* populations that left the plate (A, C) and the choices made by the worms that remained on the plate (B, D). In panel B, positive vs negative values reflect attraction to the first-vs second-listed type of bacteria, respectively. In panel D, positive vs negative values reflect attraction to the inoculated vs uninoculated half of the plate, respectively. Each large dot is the mean of 5 biological replicates (shown as transparent blue dots). Error bars represent 95% confidence intervals. ** *p* < 0.01.

**Figure 2.**
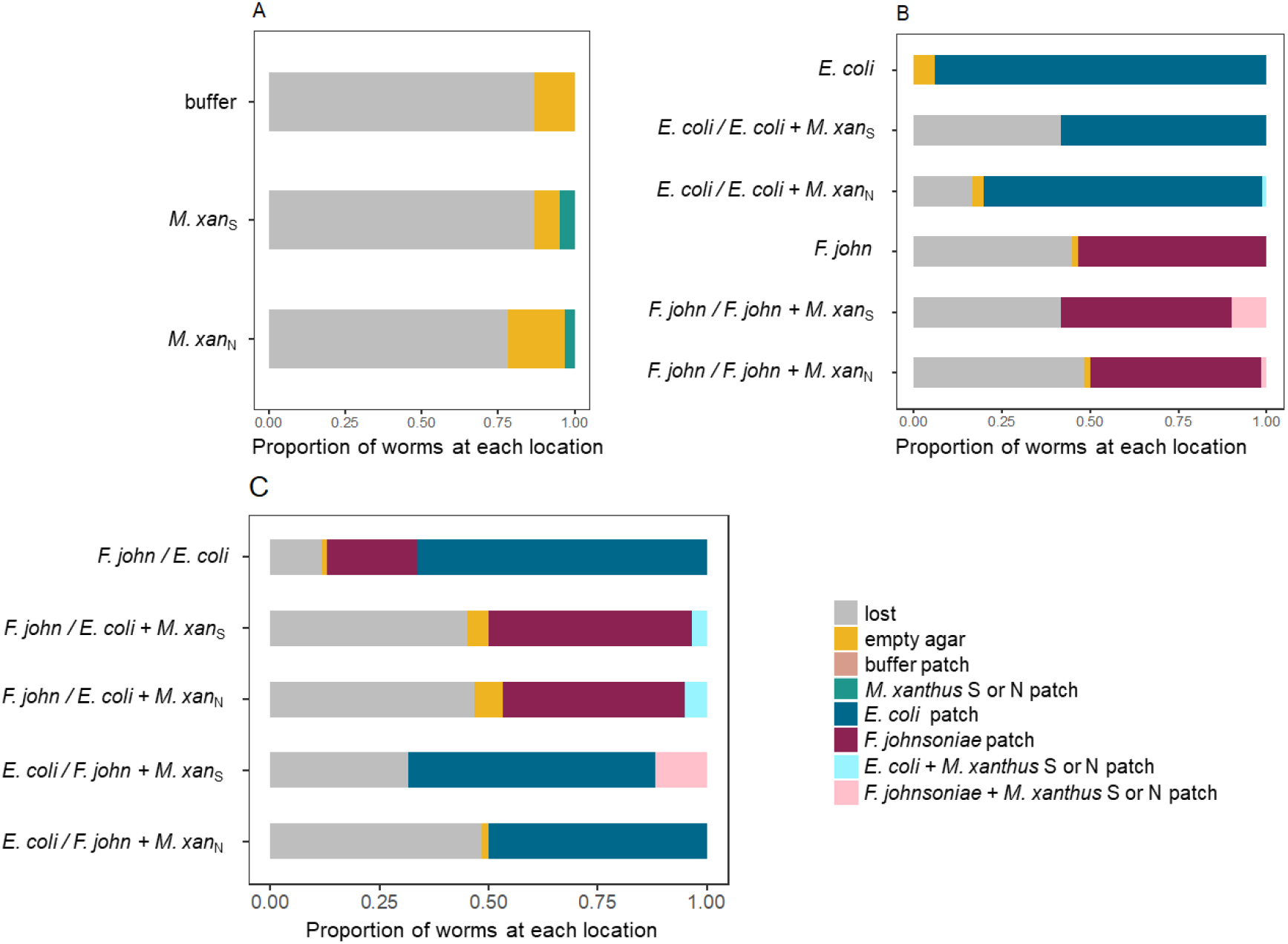
*C. elegans* prefers prey patches not containing *M. xanthus*. Here we show worm locations relative to two circular prey patches after 25 hours on plates where either (A) the patches contained either buffer or *M. xanthus*, (B) one patch contained *E. coli* or *F. johnsoniae* and the other patch contained the same with or without pre-mixed *M. xanthus*, or (C) one patch contained *E. coli* or *F. johnsoniae* and the other patch contained the other prey species with or without pre-mixed *M. xanthus*. Worms found on the plate but not in a patch are indicated in yellow, and worms that had left the plate by 25 hours are indicated in grey. Bars are mean worm counts from 3 biological replicates. ‘*F. john*’ = *F. johnsoniae*, ‘*M*.*xan*_N_’, ‘*M*.*xan*_S_’= *M. xanthus* strains N and S, respectively.

**Figure 3.**
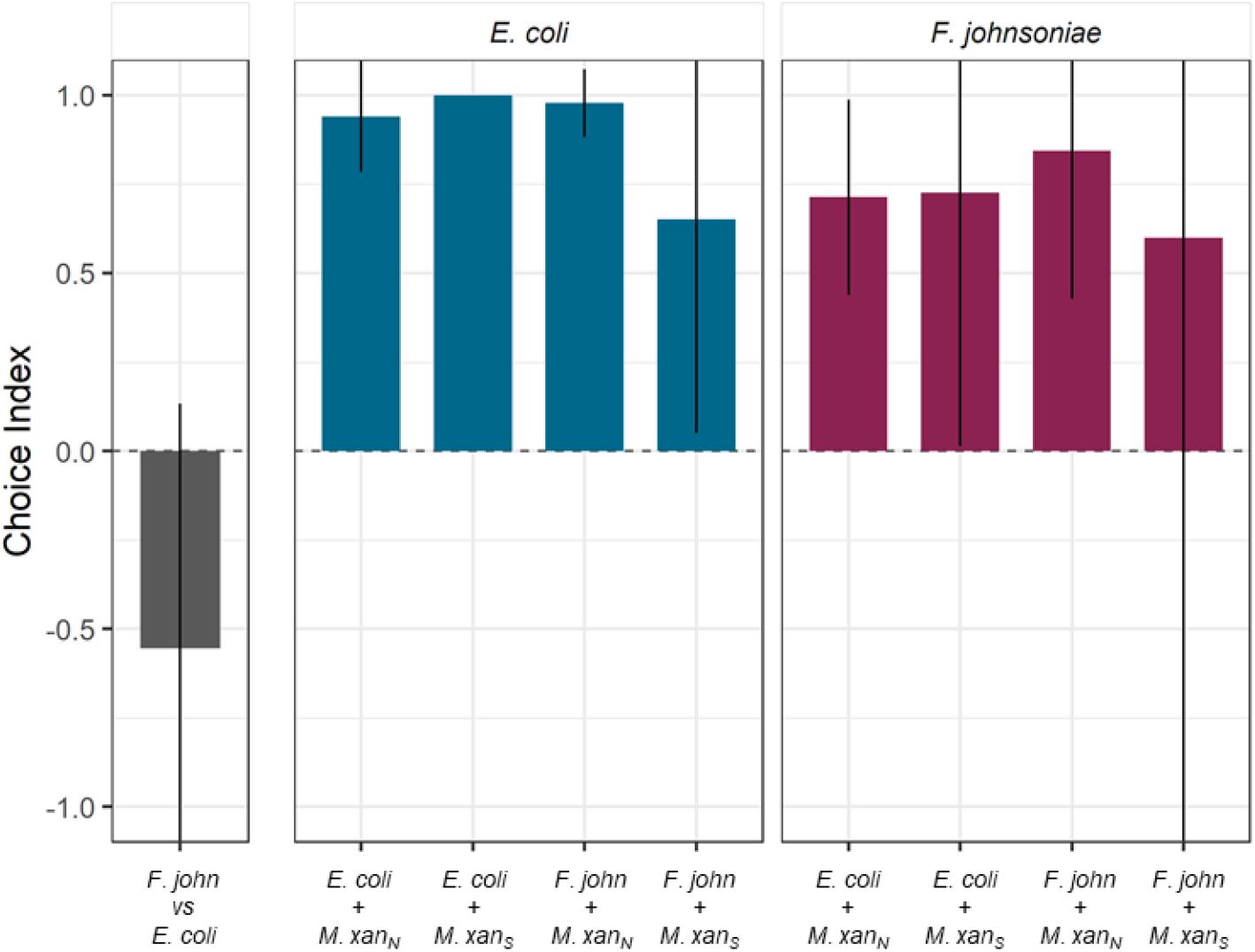
Choice indices for circular patches of basal prey. The leftmost section shows the preference of *C. elegans* for *E. coli* over *F. johnsoniae*. The next two sections show the choice of the worms between a mono-species patch of basal prey (indicated above the panel, *E. coli* = blue or *F. johnsoniae* = red) and a second patch (indicated on the x-axis) consisting of basal prey mixed with *M. xanthus* either strain S or strain N. Error bars are 95% confidence intervals. For the choice *E. coli* / *E. coli* + *M. xan*_S_ the error bar is zero. ‘*F. john*’ = *F. johnsoniae*, ‘*M*.*xan*_N_’, ‘*M*.*xan*_S_’= *M. xanthus* strains N and S, respectively.

### C. elegans half-plate choice assays

We drew center lines on CFcc plates and prepared both living and dead cultures of *M. xanthus* strains S and N as reported above. We inoculated one half of each petri dish with 100 µl of one of the bacterial cultures or buffer control spread with a 10 µm loop and allowed the inoculum to dry. We bleached *C. elegans* and adjusted the egg suspension to 50 eggs/µl by counting the number of eggs in three 0.5-µl drops. We immediately added 20 µl of egg suspension (approximately 1000 eggs) to each dish along the center line. We incubated the dishes at 25 °C and 50% rH and counted the number of worms on each side of the dish under a dissecting microscope at several time points. Results from these assays are shown in Fig. 1C,D.

### M. xanthus swarming assays

We marked CFcc plates with reference lines and scale bars for image analysis. We inoculated 15 µl each of *F. johnsoniae, E. coli*, and *M. xanthus* strain S in a row, with *M. xanthus* in the middle and 1 cm distance between each inoculation spot, and incubated the dishes at 20 °C and 50% rH. We prepared *C. elegans* worms by bleaching them on the same day that we plated the bacteria, and the next day we added the appropriate number of worms to each dish either by manually picking the desired number of individual L1 larvae and adding them directly or by adding 10 µl of worms suspended in CFcc liquid adjusted to the appropriate concentration. We took pictures of the experimental plates every 24 hours, and we measured the distance *M. xanthus* swarmed into each prey patch over time with image analysis using Fiji^61^. Results from these assays are shown in Fig. 4.

**Figure 4.**
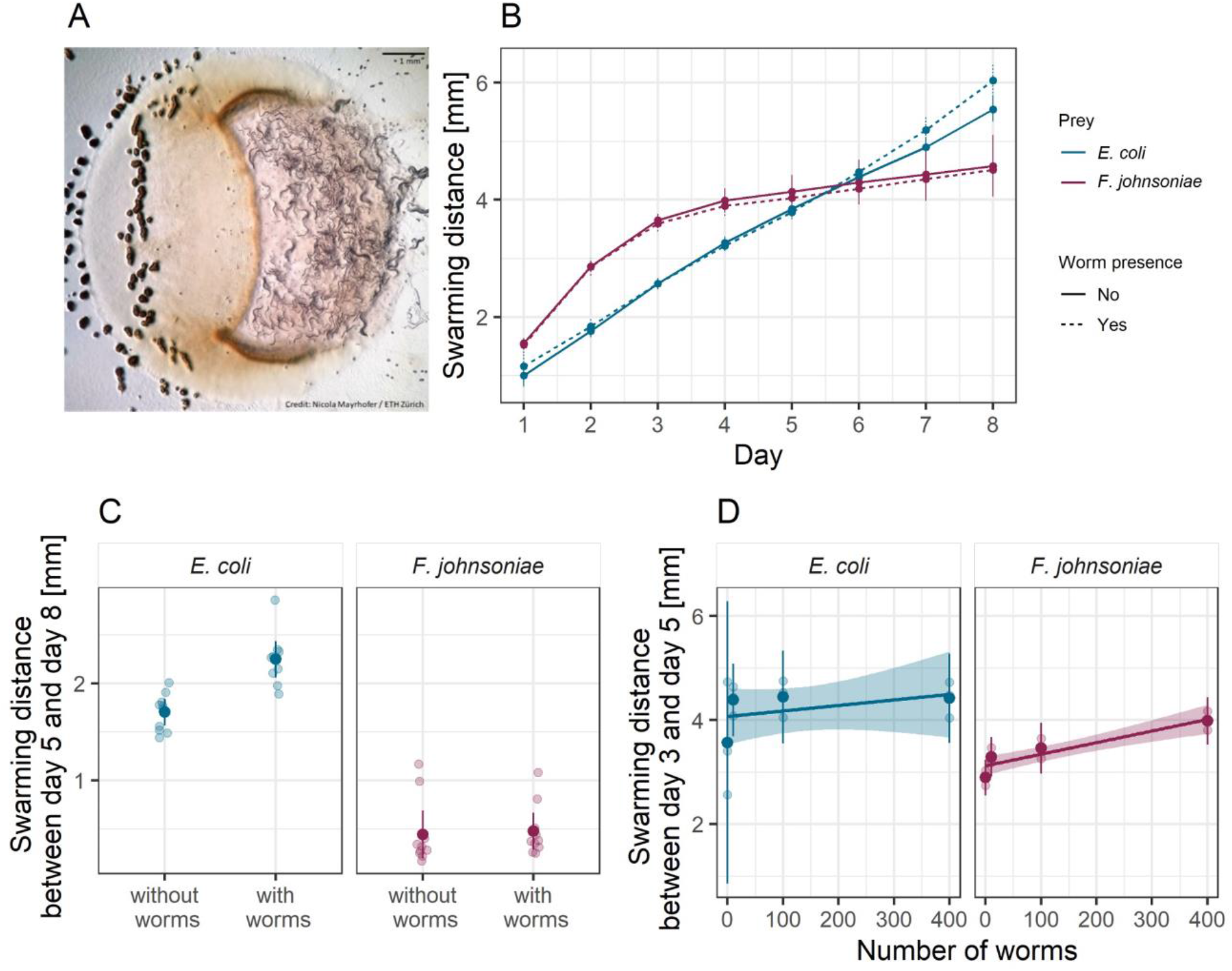
*C. elegans* presence increases *M. xanthus* predatory swarming rate. (A) Picture of *C. elegans* (right) and *M. xanthus* strain S (orange lawn and black fruiting bodies on the left) preying upon an *E. coli* patch (raised circle). We estimated the predatory performance of *M. xanthus* as the swarming distance along the midline of the prey patches of *E. coli* (blue) and *F. johnsoniae* (red). We show the swarming distance over time (B) in the presence (dotted lines) and absence (plain lines) of *C. elegans* populations initiated with ten worms (which started reproducing at day 5), and the associated total swarming distances between days 5 and 8 (C) from the same experiment. Panel (D) depicts the total swarming distances between days 3 and 5 in the presence of different numbers of *C. elegans* (the worms did not reach maturity until day 5 and so did not reproduce during this experiment). Each large dot is the mean of five biological replicates (shown as transparent dots). Error bars and shaded areas represent 95% confidence intervals of the means and the regression lines, respectively.

### Statistical analysis

We performed all data analysis and statistical testing using R version 3.6.2 and RStudio version 1.2.5033^62,63^. We tested the effect of *C. elegans* on *M. xanthus* swarming distance on prey using a mixed linear model with prey type (*E. coli* or *F. johnsoniae*) and worm treatment (factor presence/absence in one experiment, continuous variable number of worms in another) as fixed effects. As we measured the swarming distance on the two prey species from the same experimental petri dish, we included the dish identity as a random factor to account for repeated measures. We compared treatment modalities using the Tukey method for multiple comparisons from the emmeans package version 1.4.3^64^. To evaluate *C. elegans*’ choice in the binary choice and half-plate choice assays, we calculated a choice index as in Moore *et al*.^65^:

(# worms on side A - # worms on side B) / (# worms on side A + # worms on side B).

Null values indicate that the worms do not prefer one side over the other (or that they all left the dish). To compare their attraction to or avoidance of live and dead *M. xanthus* (strain S or N), we used an ANOVA with the options treatment and time as fixed effects. We considered time as a fixed effect because we were interested in whether the differences between the option treatments would change over time. We performed post-hoc comparisons with the Tukey method. We further tested whether the worms preferred one option over the others with one-sample *t*-tests against 0 with Bonferroni correction for multiple testing.

## Results

*Only a few C. elegans worms interact with M. xanthus regardless of whether it is alive or dead*. To test for interactions between *M. xanthus* and *C. elegans*, we co-cultured worms and bacteria on agar petri dishes in a variety of assays. In all of these assays, the worms had three spatial areas to choose among: i) outside the assay plate, which could be reached by worms climbing out of the petri dish, ii) agar-surface regions with no bacteria present and iii) agar-surface areas covered by bacterial cells, with some of these areas being circular patches (binary choice assays) and others covering half of a petri dish (half-plate choice assay).

In our first experiments, only *M. xanthus* was available as potential prey, and we offered both live and dead *M. xanthus* cells to *C. elegans* to test for any effect of cell death on their attractiveness. If *M. xanthus* is an attractive prey item for *C. elegans*, it might attract worms equally whether alive or dead. Alternatively, if *C. elegans* avoids living *M. xanthus* cells because they produce a repellant compound, dead *M. xanthus* might nonetheless serve as a palatable food source. Two strains of *M. xanthus* were offered to *C. elegans*, one motile (strain S) and one non-motile (strain N).

In a binary choice assay, we inoculated two circular patches of *M. xanthus* on an agar surface and added L1 larval worms to a bacteria-free region of the plate, equidistant from the two *M. xanthus* patches. In this assay, we counted how many worms left the plate vs. remained on the plate after 1 and 18 hours and, of those that remained, how many entered one or the other of the *M. xanthus* patches. In most replicates, regardless of the options provided, large majorities of the worm populations emigrated from the dish (>75% on average), and there was no general difference in the rate of emigration as a function of *M. xanthus* strain identity (ANOVA bacterial identity factor *F*_*2,12*_ = 0.8, *p* = 0.5, Fig. 1A). Those who stayed demonstrated no clear general preference between live vs dead patches across both strains and both examined time points (1 and 18 hours, ANOVA choice:time interaction *F*_*2,24*_ = 5.67, *p* < 0.01, post-hoc Tukey HSD tests *p* > 0.2; Fig. 1B). One exception to the general lack of a strong effect of *M. xanthus* death occurred on dishes containing the motility mutant strain N. On these plates, the worms seemed to initially prefer the live strain N patch after 1 h but then changed their preference to the dead patch by 18 hours (post-hoc Tukey HSD test *p* = 0.01). Because no similar pattern was seen for strain S, this result suggests that the effect of death on the attractiveness of bacterial cells to nematode predators can vary across conspecific genotypes.

The preference of the worms to leave the experimental petri dishes suggested that *M. xanthus* might repel *C. elegans*. To test this hypothesis, we inoculated *M. xanthus* alone – either strain S or N, alive or dead – onto half of the agar surface of the petri dish, leaving the other half uninoculated. In the previous binary choice assay, the patches of *M. xanthus* were small relative to the agar surface of the experimental petri dishes, reducing the likelihood of finding worms in a patch unless the bacteria actively attract them. In contrast, in this half-plate choice assay we would expect to find 50% of the worms on the plate located in the bacterial lawn, assuming no interactions between the two organisms. We included plates inoculated with sterile resuspension buffer to control for the potential attractive or repulsive effect of the buffer itself. We added *C. elegans* eggs to the midline of the dish and counted how many worms left vs remained on the plate after hatching and, of those that remained, how many went to each half of the dish.

The overwhelming majority of worms (>90%) preferred to leave the dish both in the absence and presence of *M. xanthus* (Fig. 1C). However, a larger proportion of worms remained on the plate when *M. xanthus* was present (∼4% on average across all treatments with *M. xanthus*) than on control plates with buffer alone (∼1%, *t*_*22*_ = 6.42, *p* < 0.01). Moreover, the worms that remained were more likely to be located on the inoculated side of the dish on plates with *M. xanthus* versus on plates with only buffer (ANOVA bacterial presence *F*_*2,19*_ = 27.43, *p* < 0.01, post-hoc Tukey HSD test for preference of strain S or strain N over buffer, *p*-values < 0.01; Figs. 1D, S1B). We saw no difference between live and dead (post-hoc Tukey HSD test *p* = 0.24) or motile and non-motile *M. xanthus* (post-hoc Tukey HSD test *p* = 0.44). Our results reveal intrapopulation heterogeneity in whether worms remain on plates containing only *M. xanthus* and suggest that, among the minority of worms that do remain, *C. elegans* is attracted to *M. xanthus*.

*C. elegans prefers both basal prey species over M. xanthus*. To investigate potential behavioral responses of *C. elegans* to *M. xanthus* relative to other potential prey, we performed additional binary choice assays. We inoculated two patches of bacteria on each dish, allowing the worms to choose between them, remain in the open agar, or emigrate from the dish and then counting the number of worms in each plate area after 1, 17 and 25 hrs. In these experiments, each patch contained either one basal prey species alone, one basal prey mixed 1:1 with *M. xanthus* strain S or N, or *M. xanthus* strain S or N alone. We added 20 worms to a bacteria-free region of the plate, allowing them to explore across the agar surface and seek out their preferred prey.

As expected from the previous results (Fig. 1A,1C), when *M. xanthus* was the only option, most worms had either left the dish entirely or were located in the open agar after 25 hours (Figs. 2A, S2, and S3). When *C. elegans* could choose between a patch of basal prey mixed with *M. xanthus* and a patch of the same prey without *M. xanthus*, the worms almost invariably chose the latter (ANOVA *F*_*13,28*_ = 10.01 *p* < 0.01, *p*-values < 0.05 except for *F. johnsoniae* v. *F. johnsoniae* + strain S; Fig. 2B). In the absence of *M. xanthus, C. elegans* preferred *E. coli* over *F. johnsoniae* (one-sample *t*-test for choice index < 0 *t*_*2*_ = -3.47 *p* = 0.037; Figs. 2C and 3). However, the presence of *M. xanthus* in *E. coli* patches altered the preference of the worms; *C. elegans* tended to prefer patches of *F. johnsoniae* over mixed patches containing both *E. coli* and *M. xanthus* (*p* = 0.096 and *p* = 0.028 for mixes with strain S and strain N, respectively, 14 two-sided *t*-tests with Bonferroni-Holm correction; Figs. 2C and 3). In general, the presence of *M. xanthus* drastically reduced patch attractiveness for *C. elegans*, independent of *M. xanthus* strain identity (*p* < 0.1, *t*-tests as described above; Figs. 2, 3, and S2).

*M. xanthus responds behaviorally to C. elegans in a prey-dependent manner*. To characterize potential behavioral responses of *M. xanthus* to the presence of *C. elegans*, we performed a binary choice assay similar to those above. In this assay, we added *C. elegans* to an agar petri dish inoculated with two patches of basal prey, one of *E. coli* (e.g., Fig. 4A) and one of *F. johnsoniae*, and one patch of *M. xanthus* at the midpoint between the prey such that it would encounter them upon swarming outward. Nematodes might alter *M. xanthus* swarming behavior due to direct interactions with *M. xanthus* or due to indirect effects of resource competition for basal prey, which in turn might be affected by the type of basal prey environment.

We measured *M. xanthus* swarming rate in each prey-patch type on dishes with (Fig. 4A) and without worms. This swarming-rate measure encompasses both the ability to penetrate the prey patch and predatory performance inside the patch^43^. For the treatment with *C. elegans*, we added 10 worms to a bacteria-free region of the plate. The nematodes had no effect on *M. xanthus* swarming rate in the *F. johnsoniae* patches (ANOVA basal prey identity:worm presence interaction *F*_*1,36*_ = 8.82 *p* < 0.01, post-hoc Tukey HSD test *p* = 0.99; Figs. 4B and C) but significantly increased *M. xanthus* swarming in the *E. coli* patches (post-hoc Tukey HSD test *p* = 0.0004; Figs. 4B and C).

The effect of worms in the *E. coli* patches only became visible after day 5 (Fig. 4C), and we hypothesized that this could be explained by the onset of worm reproduction, as the worms reached maturity, and subsequent increase in the worm population size. To evaluate whether larger populations of *C. elegans* lead to an increase in *M. xanthus* swarming rate, we repeated the experiment using different numbers of worms. For this experiment, we show the swarming distances between days 3 and 5 (rather than days 5-8 as in the previous assay) to capture the response of *M. xanthus* to the inoculated number of worms rather than to a growing population, as within this time frame the worms had not yet completed their development. We found that *C. elegans* increased *M. xanthus* swarming on *E. coli* to a similar small degree regardless of worm population size (linear model *F*_*1,10*_ = 0.82, *p* = 0.39, adjusted R^2^ = -0.01; Fig. 4D). It therefore remains unclear whether the time delay is simply a delay in *M. xanthus* response or whether it has to do with the developmental progress of the worms. However, in contrast to our first experiment, *C. elegans* clearly increased *M. xanthus* swarming rate on *F. johnsoniae*, but did so only as a function of worm population size (linear *F*_*1,10*_ = 32.01, *p* < 0.001, adjusted R^2^ = 0.74; Fig. 4D), such that an effect of *C. elegans* was only evident when hundreds of worms were added.

## Discussion

Despite their importance in microbial population turnover and community dynamics, there has been little research on microbial trophic chains. Here we investigated interactions between two bacterivorous predators, *M. xanthus* and *C. elegans*, in the context of a synthetic community that included two species of basal prey (*E. coli* and *F. johnsoniae*). We found that *M. xanthus* generally repels *C. elegans* relative to the effects of the two basal prey. When *M. xanthus* was the only prey option available, most worms departed our experimental predation dishes (Figs. 1A, 1C and 2A), whereas when either or both of the basal prey were offered most of the worms remained on the plates (Fig. 2B, C). *M. xanthus* was not entirely repulsive to worms; when only *M. xanthus* was offered, the few worms that remained on the plate localized more frequently within areas with *M. xanthus* than on open agar (Fig. 1D). However, the presence of *M. xanthus* in a prey patch mixed with a basal prey species repels *C. elegans* when a separate monoculture patch of either of the basal prey species is available (Figs. 2B, 2C and 3). We further showed that *C. elegans* can alter *M. xanthus* behavior by increasing the bacterial predator’s swarming rate across patches of basal prey, and that such behavior alteration depends on basal-prey identity. Our results highlight the importance of predator-predator interactions in microbial communities and the idea that other community members may often considerably modify pairwise behavioral interactions between organisms in such communities.

Theory and previous experiments in metazoans predict that the behavioral interactions between apex predators and mesopredators are driven mainly by the former^66–69^. In our microbial system, however, when a pure patch of basal prey was available, *C. elegans* avoided foraging areas occupied by *M. xanthus* even when they also contained the worms’ preferred prey (Figs. 2 and 3). Although in this study we did not formally test whether *C. elegans* can use *M. xanthus* as a food source to fuel worm population growth, our results suggest that the behavioral interactions between the two may prevent *C. elegans* from acting as an apex predator in this system. Despite Dahl and colleagues’ conclusion that *C. elegans* can be a predator of *M. xanthus*^48^, we found that *C. elegans* worms seem to consider *M. xanthus* to be an unpalatable food source. They avoid it whenever more palatable prey is available (Figs. 2B, 2C, 3), and a majority of individuals avoid it even when there is no other prey available (Figs. 1A, 1C, 2A). *C. elegans* does not similarly avoid the basal prey (Figs. 2B, 2C).

*C. elegans* uses its nervous system to recognize different bacteria in its environment^70^ and to modify its locomotive behavior in response to prey quality^71^. It can learn to recognize and approach high-quality prey^71^ and to avoid pathogens^72^. There is evidence that learned pathogen avoidance is modulated by changes in gene expression that are heritable through four generations^65^. While the mechanistic reasons for the avoidance behaviors we observed here remains to be investigated, we can hypothesize that *M. xanthus*’ large secondary metabolome^38,73–75^ contains some compounds with a primary or secondary defensive function against predators, and that the worms can either sense them at a distance or learn to avoid them after the first encounter^72^. The extent to which such compounds are repellant for the worms may be modulated by the basal prey species in the context of different multispecies setups, either because *M. xanthus* produces them only during predation or because they are repellant only compared to the more attractive compounds produced by the basal prey. If such compounds are discovered, it would be of interest to investigate whether they have specific targets or can repel a broad range of predators, to assess the importance of chemical warfare in *M. xanthus*’ trophic interactions.

While most of the *C. elegans* population avoided *M. xanthus* by leaving the experimental plates entirely, minorities of worms did interact with *M. xanthus* (Figs. 1A, 1C and 2A) and preferred it over sterile buffer (Fig. 1C,D). This could reflect a level of behavioral heterogeneity in the worm population, potentially due to different feeding preferences, predatory behaviors, or sensitivity to repellant compounds across individual worms. Regardless, there may be some conditions under which *C. elegans* can be attracted to *M. xanthus*, for example when no other prey source is available.

Still, the primary response to *M. xanthus* we observed when other prey were available was repulsion. Our results do not support the hypothesis that *M. xanthus* facultatively produces repellant compounds in response to the presence of *C. elegans*, as dead bacteria (that no longer produce any compounds) were not collectively less repellant to the worms than live bacteria (Fig. 1). Alternatively, we can hypothesize that repellant compounds produced constitutively or with a different original purpose during the growth phase of *M. xanthus* remain active, at least partially, in the inoculum and deter *C. elegans* from interacting with the bacterial predator even after it is dead. For strain N in the binary choice assay, the worms that stayed on the plate were more likely to interact with the dead bacteria after 18 hours than after only 1 hour (Fig. 1B). It is possible that after 18 hours the repellant compounds had been diluted or degraded to a level that allowed *C. elegans* to interact with the dead cells. We might expect this effect to be observed more noticeably in strain N than in strain S because strain N’s inability to swarm might result in a higher concentration of any secreted compounds in the vicinity of the living bacterial colony, creating a stronger contrast between the live and dead treatments of strain N than of strain S. In addition, on plates with live *M. xanthus*, we observed that the worms remaining on those plates that entered areas with *M. xanthus* tended to aggregate around fruiting bodies. Cells undergoing development would likely decrease active production of repellant compounds in order to devote cellular resources to the developmental process. These hypotheses merit further investigation.

The impact of apex predators on mesopredators goes beyond killing effects to include indirect behavioral changes. Some predator-induced behavior changes do not require direct contact between the predator and its prey. Animal mesopredators commonly observe traces of an apex predator (e.g. scat) and modify their foraging strategies to avoid certain areas or times of day in order to reduce their own predation risk^51,69,76,77^. Such non-lethal effects, often called risk effects, shape not only mesopredators’ behavior but also their reproduction and survival^78,79^, with cascading impacts on ecosystem structure^80^. In our model system, the presence of *C. elegans* in the arena modified the predatory behavior of *M. xanthus*, even though the worms rarely interacted directly with the *M. xanthus* swarm. This effect was modulated by the basal prey identity, suggesting that prey species may exert a potential bottom-up control on the interactions between predators^81–83^. When the basal prey was *F. johnsoniae, M. xanthus’* swarming rate on the prey patch depended on the density of worms. In contrast, *M. xanthus* swarmed faster on *E. coli* in the presence of *C. elegans* irrespective of the apex predator density (after an initial delay in the response; Fig. 4). It is possible that lower attraction of the worms to *F. johnsoniae* explains the density-dependent response in the mesopredator: when the density of *C. elegans* is low, there are often by chance only very few worms in the vicinity of *M. xanthus* when it preys on *F. johnsoniae* as opposed to when it preys on *E. coli*, which is statistically less likely as the worm density increases.

The Mesopredator Release Hypothesis (MRH, *e*.*g*.^67,84^) predicts that interference interactions between apex predators and mesopredators can have profound effects on regional ecosystem structures and large-scale biomass distribution patterns. These effects have been observed in studies of animal communities^66,85^. According to the MRH, reduction in an apex predator population liberates mesopredators both from killing effects and from the need for risk-reduction behaviors. As a result, mesopredator populations increase and individuals forage more freely, which can decimate prey populations. Interference effects between top predators and mesopredators should therefore be considered in order to understand how ecosystems are shaped. Johnke and colleagues^25^ showed that, in microbes, the combination of generalist, semi-specialist, and specialist predators can help maintain overall diversity and prevent extinction of prey species due to interference competition among the predators. However, these predators did not necessarily prey on each other, and it is not known how the addition of such effects may have altered the outcome. Such studies provide valuable insight into factors maintaining diversity in microbial communities, but they are unable to address questions about more complex trophic dynamics and, in particular, the ways in which apex predators may control populations of microbial mesopredators.

In metazoans, behavioral observation often constitutes a key source of information about indirect interactions that, as previously mentioned, can alter both food web structure and dynamics, sometimes more strongly than density-mediated effects (reviewed by Werner and Peacor^86^). Our results do not yet provide a clear picture of the factors governing interactions between *C. elegans* and *M. xanthus*, but they offer a starting point for developing model experimental systems that allow systematic behavioral observation in nematodes and bacteria. Given the crucial role of organism behavior in the structuring of metazoan food webs, we emphasize the need for microbial food web studies to investigate behavior-mediated effects as well as direct killing effects. We suggest that *C. elegans* may not be an ideal candidate for the role of apex predator, given unclarity regarding its ability to prey on *M. xanthus*, but perhaps a protozoan or another nematode such as *Pristionchus pacificus* would more readily feed on *M. xanthus*.

In microbial communities, the overlap between ecological and evolutionary timescales has generated a number of insightful studies on food web dynamics and between-predator interactions^85,87–90^. Yet most work has to date focused on density-mediated effects of interactions, and conceptual strategies for studying behaviors of predators of microbes remain scarce. Our synthetic community constitutes one step forward in that direction.

## Acknowledgments

Some strains were provided by the CGC, which is funded by NIH Office of Research Infrastructure Programs (P40 OD010440). The bleaching procedure was developed with the help of Silvan Spiri and Andrea Haag from the Group of Prof. Dr. Alex Hajnal at the Institute of Molecular Life Sciences, University of Zurich. We thank Peter Zee for preliminary discussions on working with *M. xanthus* and nematodes.

## Supplementary Information

**Figure S1.**
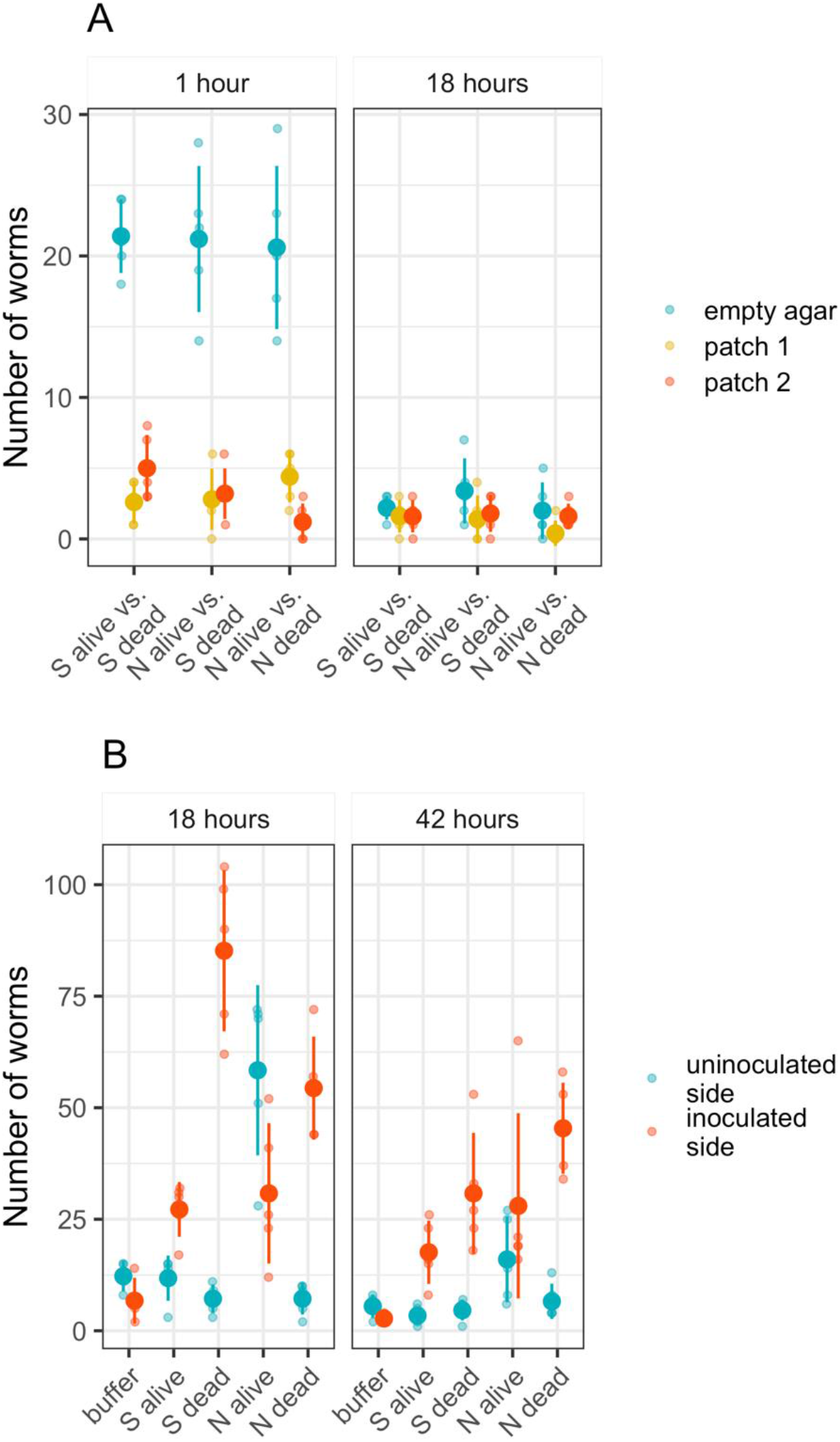
Localization of *C. elegans* on plates containing only *M. xanthus*. Numbers of worms in each specified plate location for the plates summarized in Figure 1 are shown. Worms that left the plates are not shown. Panel A shows the results of the binary choice assay with *M. xanthus* alone (Fig. 1A,B), and panel B shows the results of the half-plate choice assay (Fig. 1C,D). Large dots are the means of 3 biological replicates (shown as transparent dots) and error bars are standard deviations.

**Figure S2.**
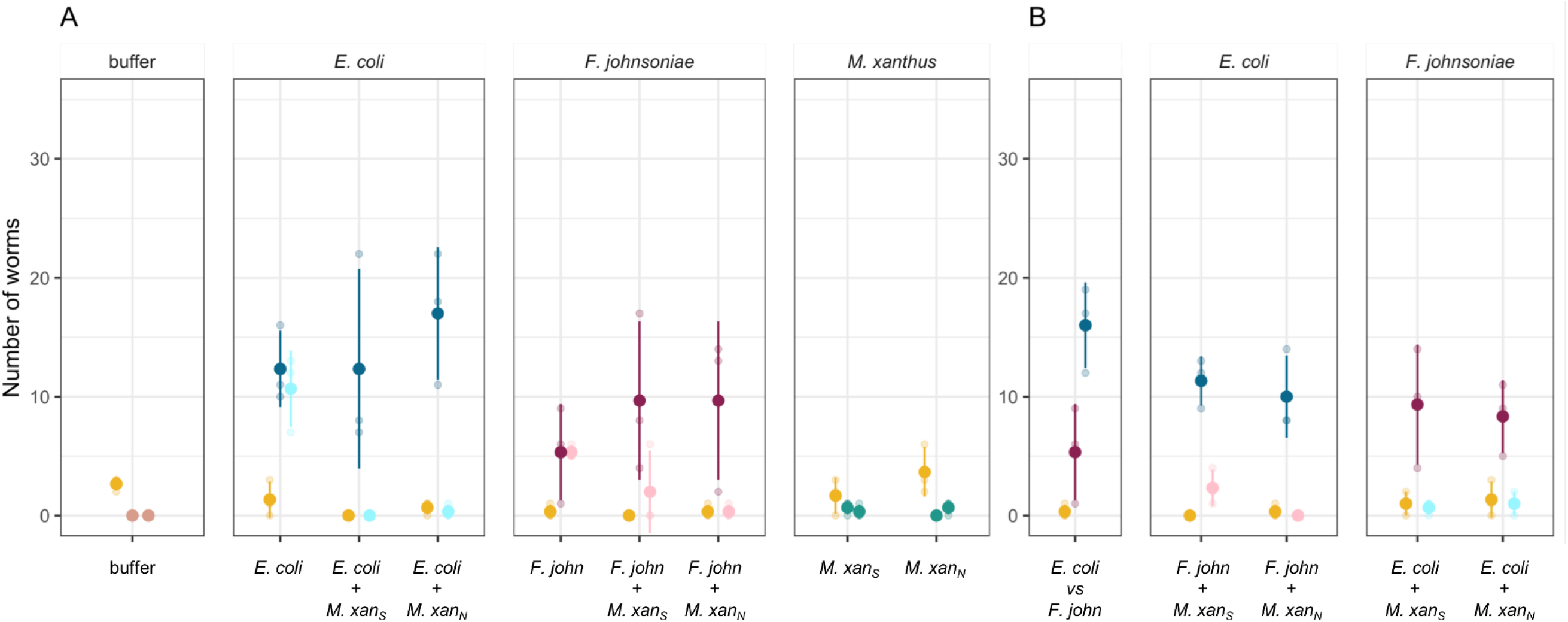
*C. elegans* prefers prey patches not containing *M. xanthus*. Here we show worm locations relative to two circular prey patches after 25 hours, on plates where both patches contained the same basal prey bacterium (A) or where the two patches contained different basal prey (B). Worms found on the plate but not in a patch are shown as yellow dots. Each separate panel shows the number of worms found in a single-species patch (or a control patch of buffer). Patch 1: buffer on the control plates (salmon dots), *E. coli* (dark blue dots), *F. johnsoniae* (dark red dots), or *M. xanthus* (green dots, left = strain S or right = strain N). Patch 2 has variously i) the same identity as Patch 1 (A), ii) the same basal prey as Patch 1 mixed with *M. xanthus* strain S or N (A; *E. coli* mixed = light blue dots, *F. johnsoniae* mixed = pink dots), or iii) the other basal prey than in Patch 1, mixed with *M. xanthus* strain S or N (B; *E. coli* mixed = light blue dots, *F. johnsoniae* mixed = light red dots). Large dots are means of 3 biological replicates (shown as transparent dots). Error bars are standard deviations. ‘*F. john*’ = *F. johnsoniae*, ‘M.xan_N_’, ‘M.xan_S_’= *M. xanthus* strains N and S, respectively.

**Figure S3.**
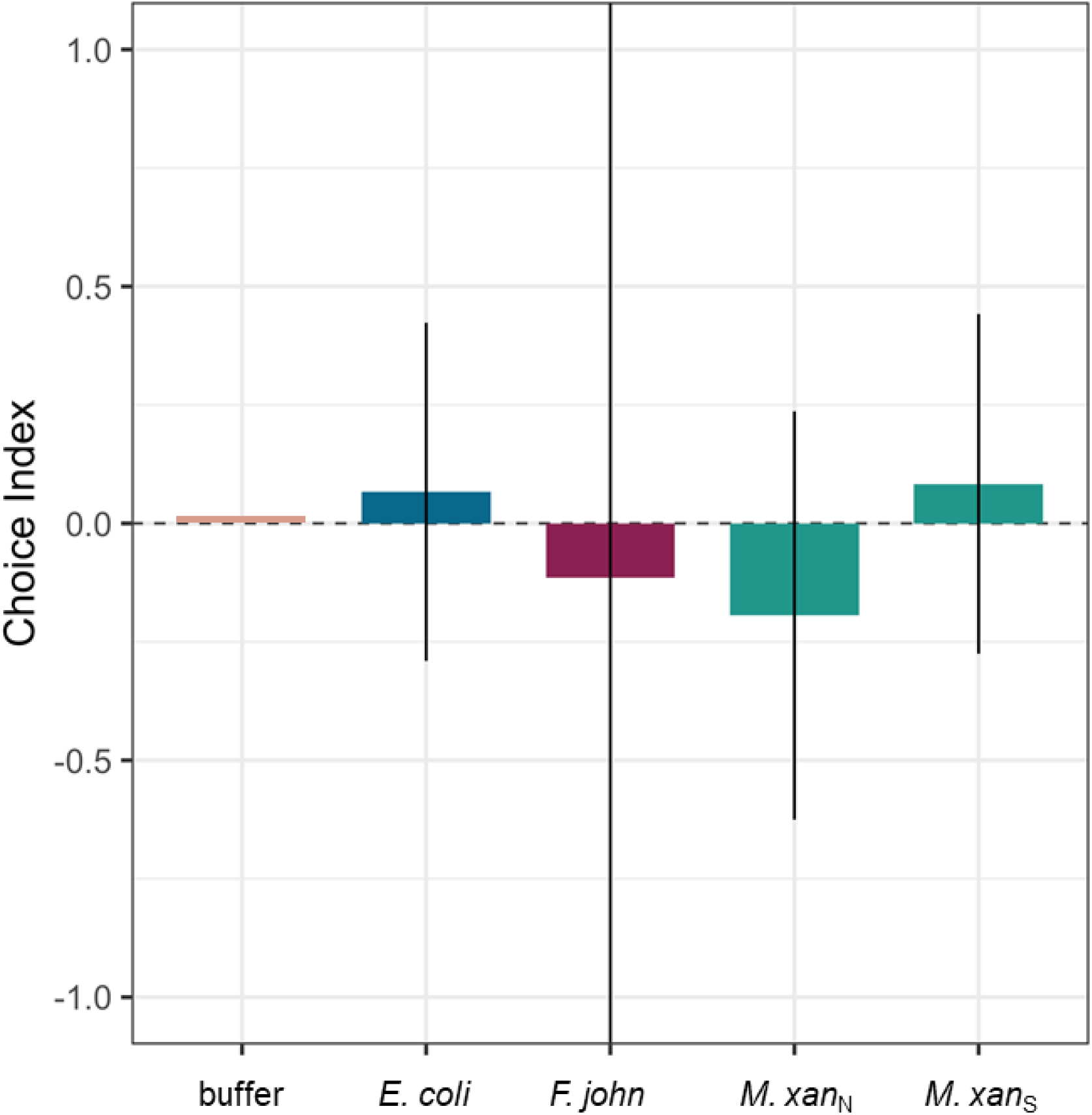
Choice indices for prey-patch controls. Here we show worm choice after 25 hours on control plates with two patches of the same prey. Colored bars show the means of 3 biological replicates and error bars are 95% confidence intervals. For the buffer treatment, the mean is zero and the error bars are zero. E. c. = *E. coli*, F. john = *F. johnsoniae*.

**Figure S4.**
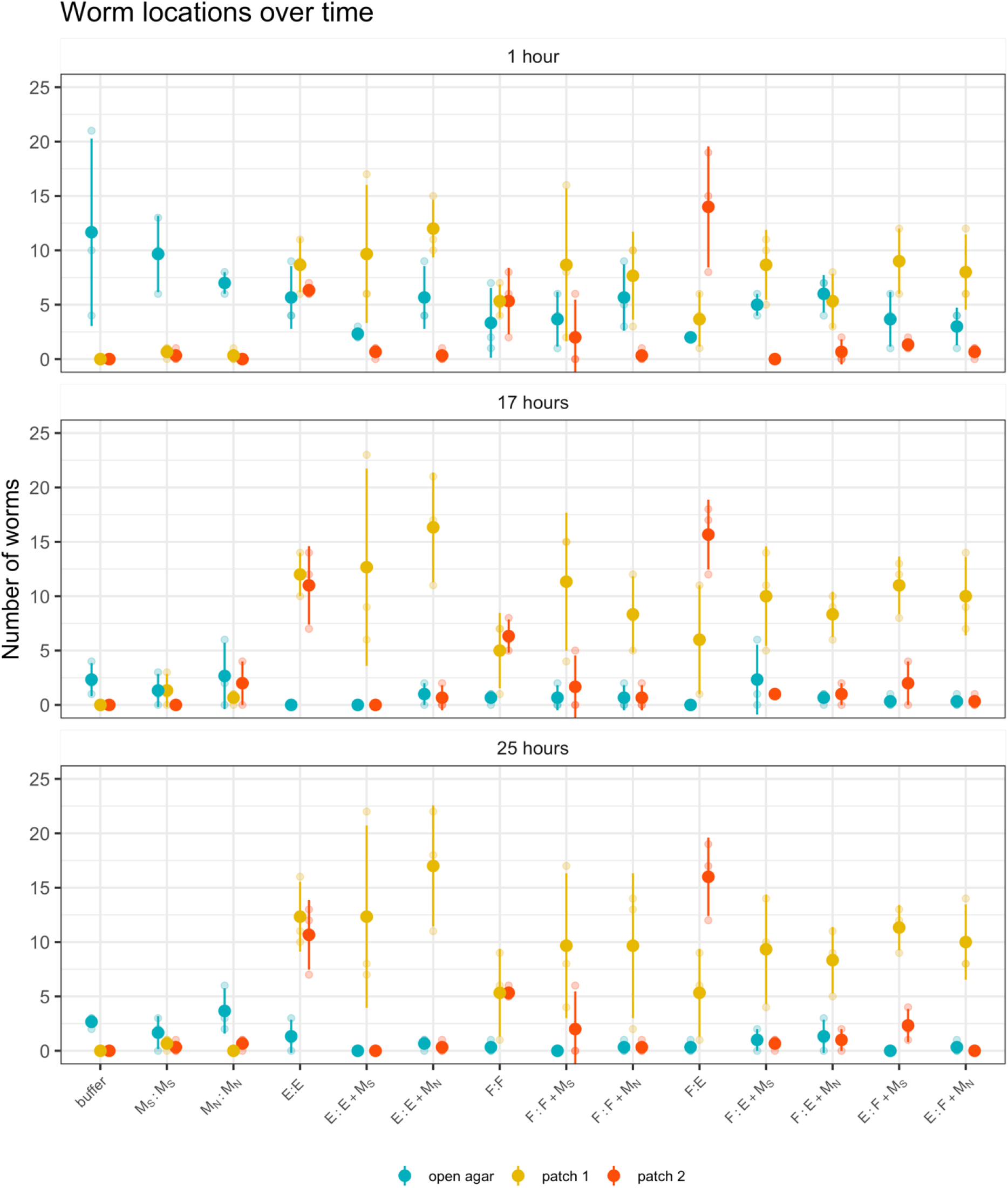
Localization of *C. elegans* on binary choice assay plates. Total number of worms in each location (either a prey patch, yellow/orange, or the open agar, turquoise) at three timepoints. Large dots are the means of 3 biological replicates (replicates shown as transparent dots) and error bars are standard deviations. E = *E. coli*, F = *F. johnsoniae*, M_S_ / M_N_ = *M. xanthus* strains S and N, respectively. For each treatment category shown on the x axis, ‘patch 1’ and ‘patch 2’ are the first and second-listed patch-identity indicators.

**Table S1.** Presence of prey bacteria in mixed patches. We tested the patches containing basal prey bacteria in the assays reported in Figure 2 to check whether basal prey bacteria could still be detected after 25 hours. The bacteria detected in each patch are shown here. Results from mixed patches where prey bacteria were still detected are indicated in bold italics. E = *E. coli*, F = *F. johnsoniae*, S = *M. xanthus* strain S, N = *M. xanthus* strain N.

| Patch 1 / Patch 2 | Patch 1 |  |  | Patch 2 |  |  |
| --- | --- | --- | --- | --- | --- | --- |
|  | Replicate 1 | Replicate 2 | Replicate 3 | Replicate 1 | Replicate 2 | Replicate 3 |
| E / E | E | E | E | E | E | E |
| F / F | F | F | F | F | F | F |
| S / S | S | S | S | S | S | S |
| N / N | N | N | N | N | N | N |
| F / E | F | F | F | E | E | E |
| E / E+S | E | E | E | S only | S only | S only |
| E / E+N | E | E | E | N only | ***E+N*** | N |
| F / F+S | F | F | F | N only | S only | ***F+S*** |
| F / F+N | F | F | F | N only | N only | ***F+N*** |
| E / F+S | E | E | E | S only | ***F+S*** | ***F+S*** |
| E / F+N | E | E | E | N only | N only | N only |
| F / E+S | F | F | F | S only | S only | S only |
| F / E+N | F | F | F | ***E+N*** | ***E+N*** | N only |

